# Genetic demultiplexing of pooled single-cell RNA-sequencing samples in cancer facilitates effective experimental design

**DOI:** 10.1101/2020.11.06.371963

**Authors:** Lukas M. Weber, Ariel A. Hippen, Peter F. Hickey, Kristofer C. Berrett, Jason Gertz, Jennifer Anne Doherty, Casey S. Greene, Stephanie C. Hicks

## Abstract

**Background:** Pooling cells from multiple biological samples prior to library preparation within the same single-cell RNA sequencing experiment provides several advantages, including lower library preparation costs and reduced unwanted technological variation, such as batch effects. Computational demultiplexing tools based on natural genetic variation between individuals provide a simple approach to demultiplex samples, which does not require complex additional experimental procedures. However, these tools have not been evaluated in cancer, where somatic variants, which could differ between cells from the same sample, may obscure the signal in natural genetic variation.

**Results:** Here, we performed *in silico* benchmark evaluations by combining raw sequencing reads from multiple single-cell samples in high-grade serous ovarian cancer, which has a high copy number burden, and lung adenocarcinoma, which has a high tumor mutational burden. Our results confirm that genetic demultiplexing tools can be effectively deployed on cancer tissue using a pooled experimental design, although high proportions of ambient RNA from cell debris reduce performance.

**Conclusions:** This strategy provides significant cost savings through pooled library preparation. To facilitate similar analyses at the experimental design phase, we provide freely accessible code and a reproducible Snakemake workflow built around the best-performing tools found in our *in silico* benchmark evaluations, available at https://github.com/lmweber/snp-dmx-cancer.

## Introduction

Sample pooling prior to library preparation is an effective strategy for experimental design in single-cell RNA sequencing (scRNA-seq) studies, which allows researchers to assess and address unwanted technological variation such as batch effects [1,2] and reduces library preparation costs [3–5]. Several strategies involve pooling cells, labeled or otherwise identifiable in some way, from multiple biological samples, followed by combined library preparation and sequencing, and computational demultiplexing to recover the sample identities of each cell. While sample pooling creates doublets consisting of cells from multiple individuals, with the doublet rate depending on the concentration of loaded cells [5], demultiplexing approaches can also identify doublets at the demultiplexing step without relying on downstream doublet identification tools [6–10]. Depending on the method used, these techniques can also avoid the phenomenon of sample index swapping, which occurs when individually prepared libraries are subsequently pooled for sequencing [11–15].

Existing demultiplexing approaches differ in their experimental procedures, computational methodology for demultiplexing, and demultiplexing accuracy. In barcoding-based approaches (e.g. MULTI-seq [16] and cell hashing [17], and GMM-Demux for doublet identification [18]), cells are experimentally tagged with universal oligonucleotides or antibodies together with sample-specific labels, which can give highly accurate demultiplexing performance. However, these approaches make sample preparation more complex, and increase costs due to reagent purchases as well as additional library preparation and sequencing. Alternatively, genetic variation-based approaches rely only on natural genetic variation between samples from different individuals (such as single nucleotide polymorphisms, SNPs), which does not require additional experimental procedures at the single-cell level. Initial genetic variation-based demultiplexing methods, such as demuxlet [5], require a known genotype reference for each sample obtained using SNP arrays, whole exome sequencing, or bulk RNA sequencing. Recently, new methods have been developed, such as Vireo [3], scSplit [4], souporcell [19], and freemuxlet [20], which can use probabilistic models to infer the genotype directly from the single-cell reads. Note that without an external genotype reference, these methods can demultiplex cells into individual samples but cannot assign cells to specific donors, since the donor identities of the inferred genotypes are arbitrary. Depending on the method, there is also the option to improve performance by providing either external sample-specific genotypes, such as from matched bulk RNA sequencing, or a list of population SNPs, such as from the 1000 Genomes Project [21] for human samples.

Recently, genetic variation-based scRNA-seq demultiplexing tools have been applied to pooled samples from cancer cell lines [22,23], using known genotype references [5,22] and pools consisting of up to dozens of cell lines. However, systematic evaluations have not yet been performed in cancer for methods that do not require a genotype reference [3,4], and using pooled samples from the same cancer type from different individuals, which are likely to be more difficult to distinguish than cell lines from distinct cancer types. Cancer is characterized by widespread additional somatic mutations, including single nucleotide variants (SNVs) [24], as well as structural variation affecting the frequency of SNVs, which could interfere with the SNP signal used to distinguish individuals in this application of demultiplexing. The frequency of additional somatic SNVs, known as the tumor mutational burden (TMB), can vary widely between cancer types [25], as well as between patients and cancer subtypes [26,27]. However, the TMB is typically small relative to the overall population SNP burden [24]. For example, population SNPs with minor allele frequency (MAF) >1% are thought to occur on the order of once per 1000 nucleotides on average, or 1000 SNPs per Mb [28]. By contrast, high-TMB cancers have been defined as having around >10 or >20 additional mutations (SNVs) per Mb [26,27] -- approximately two orders of magnitude lower frequency than the population SNPs. In the case of typical scRNA-seq protocols that sequence the 3’ end of transcripts, only SNPs within the sequenced region (e.g. 100-200 nucleotides) can be detected, but the same arguments may be applied to compare the proportion of cancer SNVs against background SNPs. Therefore, it seems reasonable to expect that the natural genetic variation signal would not be severely obscured by the TMB, and that genetic variation-based demultiplexing tools should still perform well for pooled tissue samples from the same cancer type from different individuals. However, this assumption has not been rigorously tested. Due to the finite and irreplaceable nature of tumor samples, we computationally evaluated demultiplexing algorithms to confirm that genetic variation-based demultiplexing performs adequately when applied to scRNA-seq pooling experimental designs in cancer, before committing samples to this experimental design strategy. In addition, we were interested in evaluating the degree to which these tools can reliably identify doublets consisting of cells from multiple individuals, including in experimental designs with extremely high proportions of doublets. Reliable doublet identification would allow the use of “super-loading” experimental designs, such as loading cells at very high concentration and subsequently removing identifiable doublets, providing substantial cost savings during library preparation [5,17,29]. In the future, these tools may also be well-suited for cell atlas initiatives, which are expected to cover large numbers of samples, including eventually those from cancer [30,31].

Here, we performed a benchmark evaluation of genetic variation-based demultiplexing in cancer scRNA-seq data using *in silico* simulations constructed from experimental scRNA-seq datasets with known sample identity for each cell. We evaluated two demultiplexing algorithms (Vireo [3] and demuxlet [5]) and five strategies for selecting the genotype reference list of SNPs used in the demultiplexing algorithms, including strategies that do not require a matched genotype reference. We also included varying proportions of simulated doublets by combining raw sequencing reads from multiple cell barcodes, which creates both identifiable doublets (from different individuals) and unidentifiable doublets (from the same individual). In addition, we tested performance for scenarios including a proportion of ambient RNA from simulated debris or lysed cells, by computationally assigning all reads from a percentage of cells to other randomly selected cells. In the benchmark evaluation, we considered scRNA-seq samples from two cancers that are potentially difficult to characterize: high-grade serous ovarian cancer (HGSOC) and lung adenocarcinoma. HGSOC is characterized by loss of TP53, and generally has medium to high SNV burden and high copy number variation (CNV) burden (particularly for focal copy number alterations), relative to other cancers [25,32], while lung adenocarcinoma is characterized by high SNV burden [25]. In addition, we compared against a baseline performance for healthy cells from cell lines [19]. Our results demonstrate that genetic variation-based demultiplexing provides high recall at acceptable precision-recall tradeoffs in both high CNV and high SNV cancer types, even with extremely high simulated doublet proportions. However, high proportions of ambient RNA from debris can reduce performance. Our results demonstrate that these tools support experimental designs that incorporate sample pooling. We provide a reproducible Snakemake [33] workflow based on the best-performing combination of tools for estimating a genotype reference list of SNPs and demultiplexing samples identified in our benchmark, to facilitate experimental design efforts. The Snakemake workflow is modular, allowing users to substitute alternative tools. The workflow requires a set of scRNA-seq pilot samples, access to a Linux computing cluster, some familiarity with the Linux command line, and optionally matched bulk RNA-seq samples (for the highest demultiplexing performance and doublet identification). All code for the benchmark evaluation and Snakemake workflow is freely accessible at https://github.com/lmweber/snp-dmx-cancer.

## Results

### Genetic demultiplexing in HGSOC and lung adenocarcinoma

We evaluated the performance of genetic demultiplexing algorithms for scRNA-seq samples from HGSOC (high CNV) and lung adenocarcinoma (high SNV) using a set of benchmark evaluations and Snakemake [33] workflow built around freely available tools including Cell Ranger [34], samtools [35], bcftools [36], Unix string manipulation tools (sed and awk), cellSNP [37], and Vireo [3] (**Methods** and **Figure 1**). The HGSOC samples were collected at the Huntsman Cancer Institute, and the lung adenocarcinoma dataset is a published dataset sourced from [38]. **Table 1** provides a summary of the scRNA-seq cancer datasets. Additional details on data collection and accessibility are provided in **Methods**

**Figure 1.**
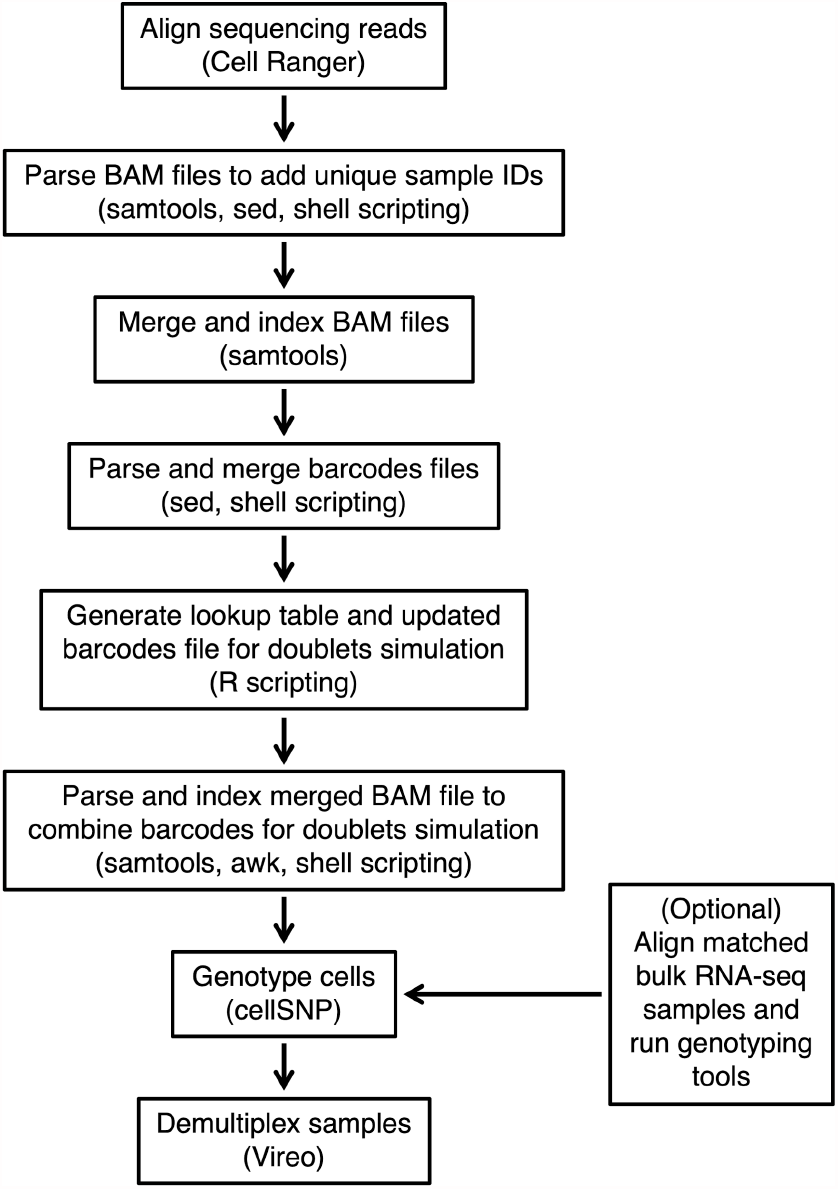
Schematic illustrating the steps in the Snakemake workflow. The workflow is designed to be modular, allowing users to substitute alternative tools. The Snakemake workflow runs a complete analysis for one dataset (HGSOC) and doublets simulation scenario (20% doublets). Our main benchmark evaluations include a second dataset (lung adenocarcinoma) and additional doublet simulation scenarios (30% doublets, no doublets)

**Table 1.**
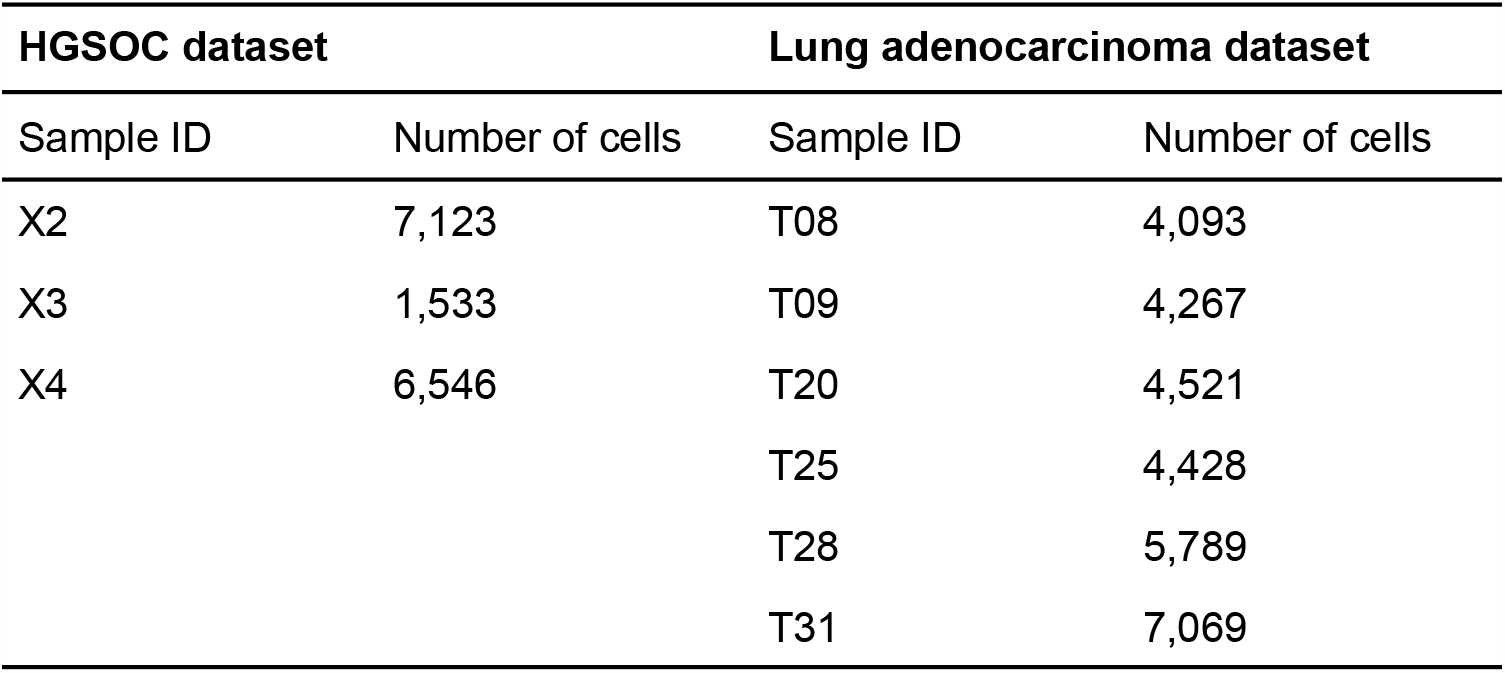
Summary of number of samples and number of cells per sample for scRNA-seq samples in HGSOC (GSE158937 and phs002262.v1.p1) and lung adenocarcinoma [38] (EGAD00001005054) datasets. The numbers of cells per sample listed are the numbers of cells provided by Cell Ranger [34] following sequencing read alignment. The HGSOC dataset additionally includes matched bulk RNA-seq samples for each sample. The lung adenocarcinoma dataset includes matched bulk whole exome sequencing samples (not used here) for each sample, but not matched bulk RNA-seq samples. Additional details for both datasets (as well as the healthy non-cancer cell line dataset from [19]) are provided in **Supplementary Table 2**

| HGSOC dataset |  | Lung adenocarcinoma dataset |  |
| --- | --- | --- | --- |
| Sample ID | Number of cells | Sample ID | Number of cells |
| X2 | 7,123 | T08 | 4,093 |
| X3 | 1,533 | T09 | 4,267 |
| X4 | 6,546 | T20 | 4,521 |
|  |  | T25 | 4,428 |
|  |  | T28 | 5,789 |
|  |  | T31 | 7,069 |

Additional and supplementary results include simulated proportions of ambient RNA from cell debris (10% and 20% debris), genotype references containing a subset of SNPs from a SNP array, and a healthy (non-cancer) cell line dataset. The optional step to run genotyping tools (e.g. on matched bulk RNA-seq samples) improved performance in our benchmark evaluations. Tools used in each step are shown in parentheses.

### High precision and recall performance using genetic demultiplexing

Using the HGSOC scRNA-seq and matched bulk RNA-seq data, we found the highest recall (defined as the proportion of true singlet cells for each sample that are identified as singlets and assigned to the correct sample) and best precision-recall tradeoff (where precision is defined as the proportion of identified cells for each sample that are true singlet cells from the correct sample), when using bcftools [36] to generate a genotype reference list of SNPs from the matched bulk RNA-seq samples, together with cellSNP/Vireo [3,37] for demultiplexing, in all simulation scenarios (no doublets, 20% doublets, or 30% doublets) (**Figure 2 a-c**). This scenario (labeled “bulkBcftools_cellSNPVireo” and colored light blue in **Figure 2**) achieves 99.0%, 99.9%, and 99.9% recall (values averaged across three scRNA-seq samples). However, in this scenario, the precision drops (100%, 85.9%, and 77.4%) (values averaged across three scRNA-seq samples) as the percentage of doublets decreases with no doublets, 20% doublets, and 30% doublets (**Figure 2 a-c**, panels from left to right), respectively.

**Figure 2.**
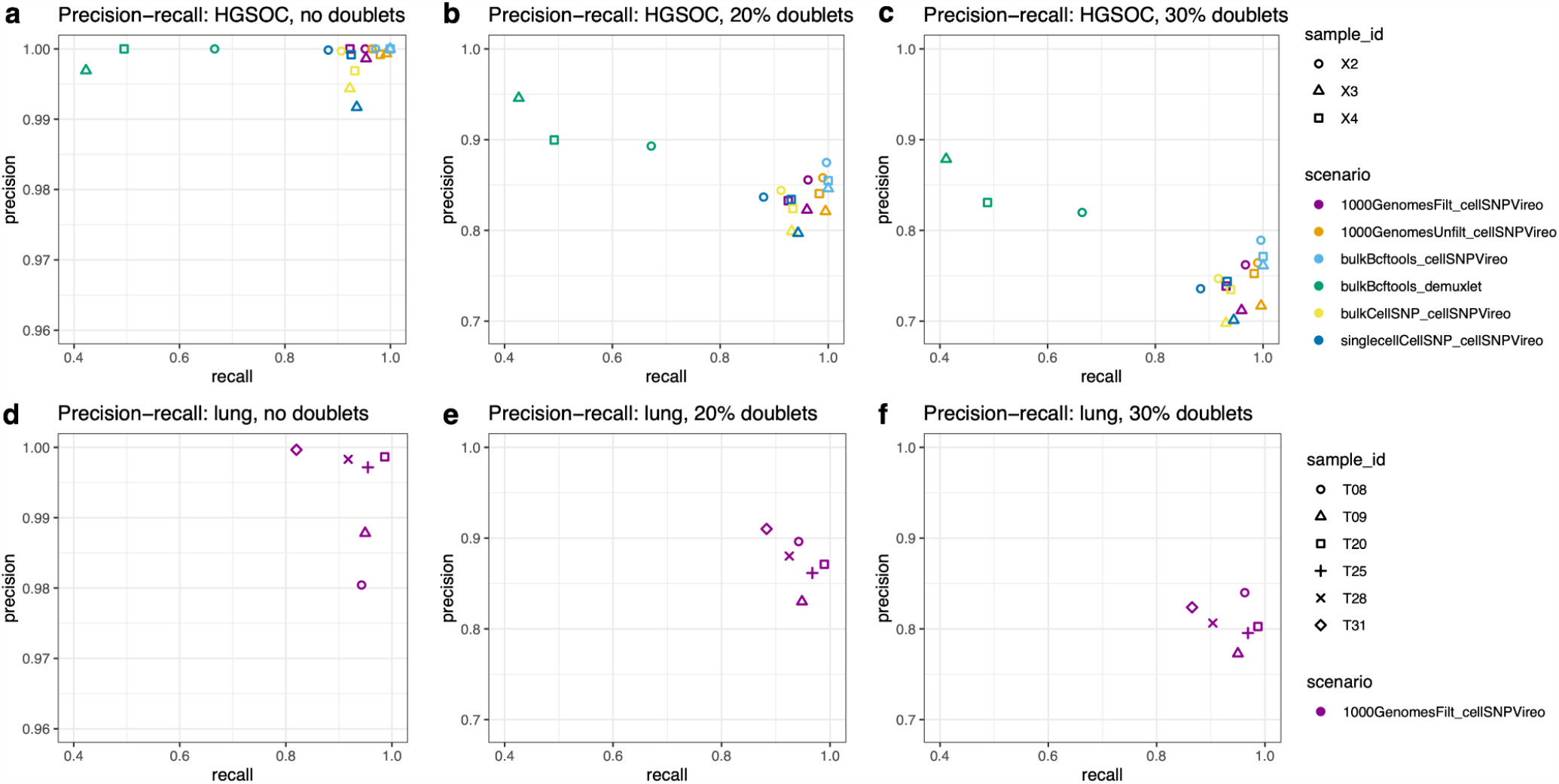
Performance evaluations for benchmark scenarios, for HGSOC dataset **(a-c)** and lung adenocarcinoma dataset **(d-f)**, across three proportions of simulated doublets (no doublets, 20% doublets, and 30% doublets). Performance is evaluated in terms of precision (y-axis) and recall (x-axis) for recovering the sample identities of true singlet cells from each scRNA-seq sample. Benchmark scenarios are labeled by color and with the naming scheme “genotypeMethod_demultiplexingMethod”. Samples within each dataset are identified with shapes. Note that y-axis limits (precision) for (a) and (d) differ from the other panels for improved visibility.

In general, we prefer higher recall at the expense of somewhat lower precision, so that we do not lose true singlet cells during the initial demultiplexing. If reduced precision is due to additional doublets that have been misclassified as singlets, these can potentially be identified and removed through downstream analyses, such as inspecting visualizations of unique molecular identifier (UMI) counts or detected genes per cell, or applying downstream doublet detection tools [6–10]. As an illustration, we investigated the types of incorrect calls leading to reduced precision in our top-performing scenario (cellSNP/Vireo with bulk RNA-seq reference) for the 30% doublets simulation in the HGSOC dataset (**Supplementary Table 3**). This showed that the doublet calls were relatively pure: of the cell barcodes called as doublets by Vireo, 99.2% were true identifiable doublets consisting of cells from distinct donors. Almost all the non-identifiable doublets (consisting of two cells from the same donor, which have the same germline SNPs) were assigned to the correct donor. By contrast, for demuxlet (HGSOC, 30% doublets, bulk RNA-seq reference), only 31.9% of the cell barcodes identified as doublets were true identifiable doublets (**Supplementary Table 4**), suggesting that cellSNP/Vireo can more reliably identify true doublets than demuxlet in these cancer samples (possibly due to non-standard allele fractions in cancer). We also applied a downstream doublet detection tool (scDblFinder [39]), but found that this did not perform well, returning large fractions of false positives and false negatives in both HGSOC and lung datasets (cellSNP/Vireo with bulk RNA-seq reference; 20% and 30% doublet scenarios) (**Supplementary Table 5**), suggesting that further analysis is required to reliably identify remaining doublets in cancer samples.

In the comparisons with demuxlet, we found that using bcftools [36] to generate a genotype reference list of SNPs from the matched bulk RNA-seq samples together with demuxlet (labeled “bulkBcftools_demuxlet” and colored green in **Figure 2**) resulted in somewhat higher precision (91.3%, 84.3%) with a large reduction to recall (53.0%, 52.1%) in the 20% and 30% doublet scenarios, respectively (**Figure 2 b-c**). However, no further improvement in precision was observed (99.9%) with a large reduction in recall (52.8%) for the no doublets scenario (**Figure 2 a**).

In the scenarios where matched bulk RNA-seq samples are not available, the next best-performing scenarios were obtained using the genotype reference from the 1000 Genomes Project [21] (provided by the authors of cellSNP/Vireo) with no filtering of SNPs (“unfiltered”) and genotype reference from the 1000 Genomes Project filtered to retain only SNPs in the 3’ untranslated region (UTR) (“filtered”, which speeds up runtime for 3’-tag sequencing protocols), together with cellSNP/Vireo for demultiplexing (labeled “1000GenomesUnfilt_cellSNPVireo” and “1000GenomesFilt_cellSNPVireo” and colored in orange and purple, respectively) (**Figure 2 a-c**). The “unfiltered” scenario achieved recall 97.9%, 99.0%, and 99.0% and precision 100%, 84.0%, and 74.5% with no doublets, 20% doublets, and 30% doublets respectively (**Figure 2 a-c**). Surprisingly, there is only a minor loss in performance in the “filtered” scenario, which achieved recall 94.3%, 95.0%, and 95.3% and precision 100%, 83.7%, and 73.8% respectively. Alternatively, when we evaluated the scenario to call SNPs directly from the scRNA-seq samples and use cellSNP/Vireo for demultiplexing (labeled “singlecellCellSNP_cellSNPVireo” and colored in dark blue), we found comparable recall (91.5%, 91.9%, and 92.1%) with a slight loss in precision (99.7%, 82.3%, and 72.7%) as the percentage of doublets decreases with no doublets, 20% doublets, and 30% doublets respectively (**Figure 2 a-c**).

Using the high-TMB lung adenocarcinoma scRNA-seq (without matched bulk RNA-seq) dataset, we only considered the scenario using the genotype reference from the 1000 Genomes Project (filtered) together with cellSNP/Vireo (labeled “1000GenomesFilt_cellSNPVireo” and colored in purple in **Figure 2**), as this resulted in the highest precision and recall in the HGSOC evaluation when using either the genotype reference from 1000 Genomes Project or directly calling SNPs from the scRNA-seq samples, while also keeping runtimes lower (details in **Figure 4**) than the 1000 Genomes (unfiltered) scenario. In this scenario (labeled “1000GenomesFilt_cellSNPVireo”), we found comparable ranges of precision and recall values as for the matching scenario in the HGSOC dataset (**Figure 2 d-f**). These results demonstrate that we can also achieve excellent demultiplexing performance even in a higher-TMB cancer setting.

### Reduced recall performance due to ambient RNA from simulated cell debris

Single-cell samples may contain proportions of ambient RNA from cell debris and lysed cells, with increased proportions in complex or necrotic samples [40], such as in cancer. Since SNPs from the ambient RNA may interfere with the set of SNPs observed for each droplet (cell barcode), this may affect SNP-based demultiplexing performance. To evaluate this effect in cancer samples, we created additional simulations where either 10%, 20%, or 40% of final cell barcodes were assumed to represent debris or lysed cells, and assigned all sequencing reads from these cells randomly to other cell barcodes (**Figure 3** and **Supplementary Figures 1-2**). The range of debris proportions was selected to be in the higher range of previously published results for non-cancer cell line data [19]. For these simulations, we included the top-performing and computationally efficient scenarios from the main benchmark (cellSNP/Vireo with bulk RNA-seq and 1000 Genomes filtered references), as well as demuxlet for comparison (HGSOC dataset). These results showed a reduction in recall performance, although the reduction was smallest for the top-performing scenario (“bulkBcftools_cellSNPVireo”, light blue). For the 10% debris simulations (**Figure 3**), we observed average recall of 85.1%, 86.2%, and 86.7% across samples (no doublets, 20% doublets, and 30% doublets scenarios) when using the bulk RNA-seq reference, and 61.7%, 60.3%, and 60.8% respectively when using the 1000 Genomes filtered reference, for the HGSOC dataset. For the lung dataset, recall decreased to an average of 57.0%, 54.2%, and 59.4% across samples (no doublets, 20% doublets, and 30% doublets) using the 1000 Genomes filtered reference. The performance of demuxlet dropped substantially, with average recall of 7.8%, 6.7%, and 6.4% across samples (no doublets, 20% doublets, and 30% doublets) using the bulk RNA-seq reference for the HGSOC dataset, suggesting that demuxlet is more sensitive to ambient RNA than cellSNP/Vireo. Additional results with higher proportions of simulated debris (20% or 40% of final cell barcodes, **Supplementary Figures 1-2**) showed greater reductions in recall performance. While precision performance was also somewhat reduced compared to the main results, the effect was much smaller than for recall (**Figure 3** and **Supplementary Figures 1-2**). Overall, these results demonstrate that ambient RNA reduces demultiplexing performance, although the effect is minimized when using the top-performing set of tools (“bulkBcftools_cellSNPVireo”, i.e. cellSNP/Vireo with a bulk RNA-seq reference, when this is available).

**Figure 3.**
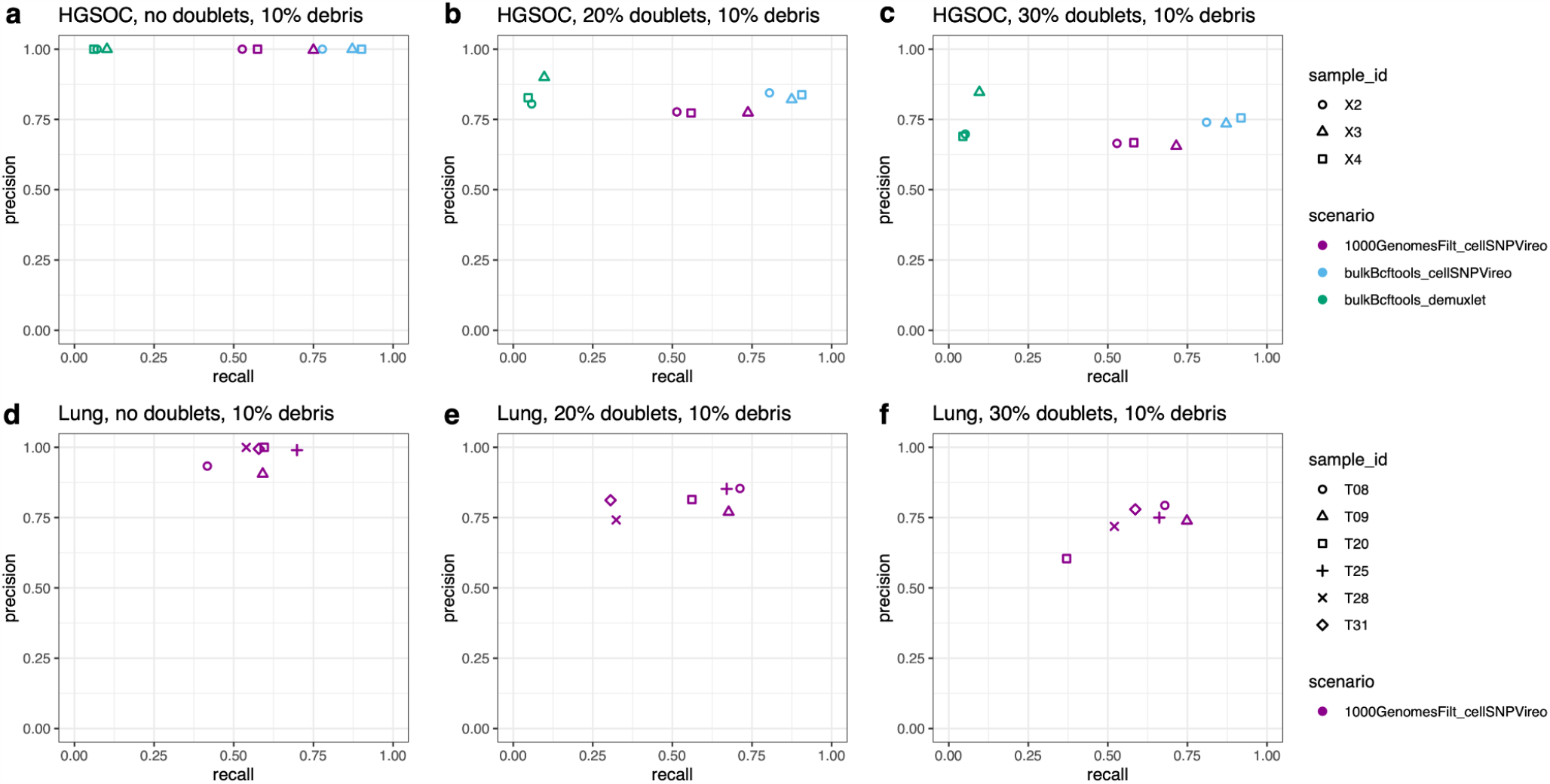
Performance evaluations for benchmark scenarios including ambient RNA from simulated cell debris. Top-performing and computationally efficient scenarios for HGSOC dataset **(a-c)** and lung adenocarcinoma dataset **(d-f)**, across three proportions of simulated doublets (no doublets, 20% doublets, 30% doublets), after introducing ambient RNA from simulated cell debris by assigning all reads from 10% of final cell barcodes to other randomly selected cell barcodes. Performance is evaluated in terms of precision (y-axis) and recall (x-axis) for recovering the sample identities of true singlet cells from each scRNA-seq sample. Benchmark scenarios are labeled by color and with the naming scheme “genotypeMethod_demultiplexingMethod”. Samples within each dataset are identified with shapes. Axis limits range from 0 to 1 for all panels.

### Performance remains high when using subset of SNPs from SNP array

Since SNP arrays are often used for genotyping in cancer studies, we were also interested in demultiplexing performance when using SNPs from a SNP array, instead of the bulk RNA-seq or 1000 Genomes references used in the main results. To test expected performance when using SNP arrays, we generated additional simulations using a subset of SNPs from a common SNP array that also overlapped with our other genotype references. We selected the Infinium Multi-Ethnic Global-8 v1.0 array from the Multi-Ethnic Genotyping Array (MEGA) consortium, calculated the overlapping sets of SNPs from this array with either the 1000 Genomes filtered (“1000GenomesFiltMEGA”), 1000 Genomes unfiltered (“1000GenomesUnfiltMEGA”), or bulk RNA-seq (“bulkBcftoolsMEGA”) reference (for the HGSOC dataset). This left either 16.5% of the 84,853 filtered 1000 Genomes SNPs, 8.6% of the 7,414,539 unfiltered 1000 Genomes SNPs, or 7.6% of the 605,367 bulk RNA-seq SNPs, or alternatively (compared to the array) 0.8%, 36.8%, or 2.6% of the original 1,733,345 array SNPs respectively (see **Supplementary Table 1** for a summary of the overlapping set sizes). Despite the large reduction in number of SNPs used for demultiplexing, performance for cellSNP/Vireo remained remarkably high, with almost no reduction in recall or precision performance in the top-performing scenario when using the overlapping SNPs from the bulk RNA-seq reference (“bulkBcftoolsMEGA”) for the HGSOC dataset (average recall across samples 96.9%, 98,7%, and 99.0%, and average precision across samples 100%, 86.3%, and 77.7%, with no doublets, 20% doublets, and 30% doublets respectively). When using the 1000 Genomes filtered or 1000 Genomes unfiltered references (“1000GenomesFiltMEGA” or “1000GenomesUnfiltMEGA”), there was a somewhat larger reduction in recall. By contrast, the performance of demuxlet was substantially lower, suggesting that demuxlet is more sensitive to the set of SNPs used for demultiplexing (**Supplementary Figure 3**). Since the set of SNPs used in the top-performing scenario (“bulkBcftoolsMEGA_cellSNPVireo”) is much smaller than the full set of SNPs from the array, and performance is only slightly reduced, these results suggest that demultiplexing performance with cellSNP/Vireo is likely to remain high when using the full array.

### High performance in baseline comparison for non-cancer cell lines

As a baseline comparison for healthy (non-cancer) data, we evaluated performance in a dataset consisting of 5 samples of induced pluripotent stem cell (iPSC) cell lines from the Human iPSC Initiative (HipSci), which was previously published by [19]. This dataset contained an average of around 9,000 cells per sample, with relatively high unique molecular identifier (UMI) counts per cell (**Supplementary Table 2**). We generated simulation scenarios containing no doublets, 20% doublets, and 30% doublets, and evaluated demultiplexing performance using cellSNP/Vireo with the 1000 Genomes 3’ UTRs filtered genotype reference. We observed similar demultiplexing performance in terms of precision and recall as in the main results for the corresponding scenarios (**Supplementary Figure 4**). These results provide a baseline comparison confirming that these demultiplexing tools perform well in non-cancer data, which is consistent with previous published results [3,19], as well as a confirmation that our simulation framework can be successfully applied in both cancer and non-cancer settings.

### Computational runtime of genetic demultiplexing workflow steps and genotyping tools

We evaluated the computational runtimes for the various components in our benchmark scenarios and Snakemake workflow using the HGSOC data. First, we found the computational runtimes for the various steps in the genetic demultiplexing workflow vary across multiple orders of magnitude and depended on whether the tool could be parallelized. The parallelizable tools (Cell Ranger and cellSNP) were run using 10 processor cores to decrease runtime, while the remaining tools used a single core. All evaluations of runtimes were performed on a high-performance Linux computing cluster. In the Snakemake workflow (**Figure 4 a**), the slowest steps were running Cell Ranger (approximately 6 hours per sample using 10 cores) and parsing the merged BAM file containing aligned reads to combine cell barcodes into simulated doublets (approximately 1 day). For the cellSNP step in the workflow, runtime depended on the choice of genotype reference list of SNPs (**Figure 4 b**). In particular, filtering the genotype reference from the 1000 Genomes Project [21] (provided by the authors of cellSNP/Vireo) to retain only SNPs in the 3’ UTRs reduced runtime from approximately 2.5 hours to less than 10 minutes (“1000GenomesUnfilt_cellSNP”vs. “1000GenomesFilt_cellSNP”), at the cost of only a small drop in performance (**Figure 2**). The runtime shown for the cellSNP step in **Figure 4 a** corresponds to the highest-performing scenario from **Figure 2** (“bulkBcftools_cellSNP”).

**Figure 4.**
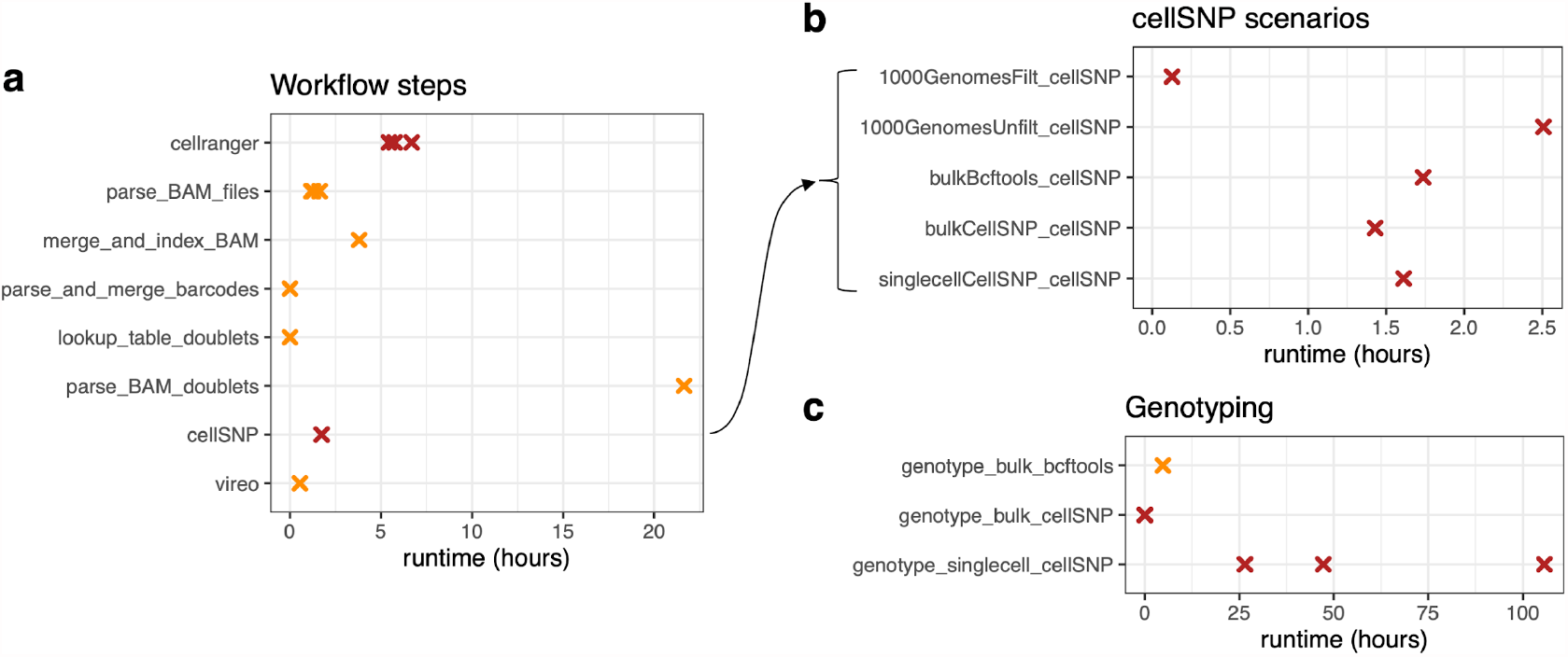
Computational runtimes (in hours) of genetic demultiplexing workflow steps and genotyping tools. **(a)** Runtimes for steps in the complete Snakemake workflow, for a single dataset (HGSOC) and doublets simulation scenario (20%). Parallelized tools (Cell Ranger and cellSNP; points indicated in dark red) were run using 10 processor cores, and all other tools using a single core (points indicated in orange), on a high-performance Linux computing cluster. For steps where samples were processed individually, separate points are shown for each sample. **(b)** Runtimes for alternative options for running cellSNP in the workflow, depending on the choice of genotype reference (1000 Genomes filtered, 1000 Genomes unfiltered, matched bulk RNA-seq using bcftools, matched bulk RNA-seq using cellSNP, and single-cell RNA-seq using cellSNP). The cellSNP step in (a) matches the row “bulkBcftools_cellSNP” in (b), which was the highest-performing scenario from Figure 2. **(c)** Runtimes for alternative options to generate genotype reference file. Horizontal axis scales differ between panels for improved visibility.

We also evaluated computational runtimes for the genotyping tools used to generate the genotype reference lists of SNPs from either the matched bulk RNA-seq samples or directly from the scRNA-seq samples (**Figure 4 c**). Here, we found by far the slowest option was to use cellSNP to generate the genotype reference directly from the scRNA-seq samples (between 1 and 4.5 days per sample using 10 cores), while generating the genotype reference from the bulk RNA-seq samples took either approximately 2 minutes per sample using cellSNP (10 cores) or 5 hours using bcftools.

## Discussion

Pooled single-cell experimental designs before library preparation together with genetic variation-based computational sample demultiplexing are a convenient and effective strategy for reducing library preparation costs and potential batch effects in scRNA-seq studies. Compared to barcoding-based approaches such as MULTI-seq [16] and cell hashing [17], demultiplexing performance may be lower depending on the quality of the genotype reference used, however genetic variation-based methods provide significant advantages in terms of simpler sample preparation and cost savings during library preparation. Here, we performed an *in silico* benchmark evaluation based on real scRNA-seq tumor tissue datasets to confirm that these tools can be effectively applied to pooled cancer samples from different individuals. We compared two demultiplexing tools (Vireo [3] and demuxlet [5]) and five genotype references. We selected HGSOC and lung adenocarcinoma, two cancer types characterized by a relatively high TMB. Previous benchmark evaluations [3,4,19] have only evaluated these tools in non-cancer datasets, which are not affected by additional mutational SNV burden that could potentially obscure the natural genetic variation SNP signal used to distinguish individuals, while previous evaluations in cancer [22,23] have relied on matched genotype references and focused on cancer cell lines, which are likely to be easier to distinguish than samples of the same cancer type from different individuals. Our benchmark evaluations include high proportions of simulated doublets (up to 30%), confirming that these tools can be used to identify singlet cells in “super-loading” experimental designs to achieve considerable cost savings in library preparation [5,17,29]. Additional analyses showed that performance remained high when using subsets of SNPs from a SNP array for the genotype reference, as well as in a baseline comparison using a healthy (non-cancer) cell line dataset [19]. However, introducing proportions of ambient RNA from simulated cell debris or lysed cells reduced performance in terms of recall, although this effect was minimized when using the matched bulk RNA-seq genotype reference. This suggests that, in cancer samples with significant proportions of ambient RNA, e.g. from cell debris due to necrosis, it may be important to consider applying experimental techniques such as straining to remove cell debris. As an illustration of expected cost savings due to lower library preparation costs in a multiplexed experimental design, we estimated library preparation and sequencing costs for designs with 4 to 8 samples, using the “Cost Per Cell” online calculator provided by the Satija Lab [29] (**Supplementary Figure 4**). We assumed 4,000 desired cells per sample after demultiplexing, i.e. after discarding identifiable doublets consisting of cells from multiple samples, but including the smaller number of non-identifiable doublets (multiple cells from the same sample, which have the same SNP profiles and cannot be distinguished using genetic demultiplexing). These designs result in cost savings of approximately 60% of the estimated cost for the experiment when using full multiplexing (all samples prepared as a single library and sequenced together) compared to no multiplexing (**Supplementary Figure 4**).

In our HGSOC dataset, we achieved the best demultiplexing performance (and relatively efficient runtimes) when using matched bulk RNA-seq samples to generate a genotype reference list of SNPs using bcftools [36], together with cellSNP/Vireo [3,37] for demultiplexing. However, using a standard list of population SNPs from the 1000 Genomes Project [21] (which does not require matched bulk RNA-seq samples) provided by the authors of cellSNP/Vireo also achieved good performance. In this case, filtering the population SNPs to retain only SNPs in the 3’ UTRs significantly reduced runtime, at the cost of only slightly lower demultiplexing performance. For the lung adenocarcinoma dataset, performance was comparable to the matching scenario in the HGSOC dataset, confirming that performance was not seriously affected by the higher TMB, and that genetic demultiplexing can be effectively applied in this setting. Since most other cancer types have lower TMB [25], we expect these results to apply to most cancer types. We provide a freely available, modular Snakemake [33] workflow implementing the best-performing scenario from our benchmark, built around cellSNP/Vireo [3,37] and other freely accessible tools, as well as additional R and shell scripts to reproduce all analyses in our benchmark evaluations and additional analyses (https://github.com/lmweber/snp-dmx-cancer), to allow other researchers to perform similar analyses for experimental design, planning, and budgeting purposes in their own datasets.

Our study has several limitations. While the best-performing benchmark scenario achieves excellent recall, precision is somewhat lower. While most doublet calls from cellSNP/Vireo in this scenario were true identifiable doublets, additional true identifiable doublets were incorrectly called as singlets, reducing precision for each demultiplexed sample. Although downstream doublet detection tools [6–10] could be applied to remove any remaining doublets, we found that this did not perform well in an initial analysis. Further work could consider a systematic evaluation of downstream doublet detection tools in the context of cancer, to complement previous results in non-cancer data [41]. In this study, we have built our Snakemake workflow around the best-performing tools (cellSNP/Vireo [3,37] for demultiplexing and using matched bulk RNA-seq samples for genotyping) and compared against demuxlet [5] and baseline scenarios (no doublets), but we have not performed a comprehensive benchmark evaluation of all available tools, such as additional tools for demultiplexing (e.g. scSplit [4], souporcell [19], and freemuxlet [20]). However, we have implemented the Snakemake workflow to be modular, so that other users may substitute alternative tools if they prefer. We also investigated the use of salmon alevin [42] for pseudoalignment of scRNA-seq reads (instead of Cell Ranger), but found that this was not compatible with the demultiplexing tools since pseudoalignment occurs at the transcriptomic instead of genomic level. However, future developments may enable conversion between transcriptomic and genomic aligned reads, and we have included alternative code scripts for salmon alevin within our code repository. Our evaluations considered only two tumor types (HGSOC and lung adenocarcinoma), and performance may differ for other cancer types or tissues. However, since we were able to demonstrate good performance in lung adenocarcinoma, one of the highest TMB cancers, we anticipate these results will also be applicable for other cancer types, which will generally have lower TMB. For the lung adenocarcinoma dataset, matched bulk whole exome sequencing data were also available for these six samples, which could be used to further improve performance using additional genotyping tools. Future work could also consider generating additional experimental data to further benchmark these tools, instead of relying on *in silico* evaluations, although in this case it may be difficult to generate a reliable ground truth.

More fundamentally, due to the reliance on genetically distinct SNP profiles, genetic demultiplexing tools work well for human samples from unrelated individuals, but performance is expected to decrease for genetically similar samples such as hereditary related human populations [19] or inbred mice, and these methods are not applicable to samples from the same individual [3]. Without a sample-specific genotype reference, it is also not possible to assign cells to specific donors, since the donor identities of inferred genotypes are arbitrary. Similarly, genetic demultiplexing does not allow identifying doublets consisting of cells from the same individual, although these are only a subset of total doublets, and decrease as a proportion of total doublets with increasing number of multiplexed samples. We also have not considered the question of identifying doublets consisting of distinct cell types (from either the same or different individuals), which may be identified using downstream analysis tools. For some experiments, a useful design strategy may also be to combine genetic-based and barcoding-based multiplexing, e.g. multiple treatments on samples from the same individual. Our Snakemake workflow can be used to demultiplex up to around 12 pooled samples without a genotype reference (limited by the demultiplexing algorithm Vireo) -- beyond this, the demultiplexing performance of the Vireo algorithm has been shown to decrease [3]. For larger experiments, if matched bulk RNA-seq samples are not available, multiple sample pools could be used, with demultiplexing done separately for each pool [3]. Splitting an experiment across multiple pools and demultiplexing within each pool also represents an opportunity to implement improved experimental designs to reduce batch effects and confounding. Finally, the Snakemake workflow is relatively computationally intensive, and requires access to a high-performance Linux computing cluster or server.

## Methods

### Benchmark evaluations and workflow

We begin by describing in detail our benchmark evaluation framework, and note that our additional Snakemake [33] workflow is built around the combination of tools that resulted in the best performance from the benchmark evaluation. Specifically, the benchmark and workflow make use of several freely available tools, including Cell Ranger [34], samtools [35], bcftools [36], Unix string manipulation tools (sed and awk), cellSNP [37], and Vireo [3]. The Snakemake workflow is designed to be modular, allowing other alternative or new tools to be substituted. All code for the benchmark evaluation and Snakemake workflow is freely available at https://github.com/lmweber/snp-dmx-cancer.

In our benchmark evaluation, we considered two genetic demultiplexing algorithms: (i) Vireo [3] together with cellSNP [37], and (ii) demuxlet [5] as an alternative genetic-based demultiplexing tool. We evaluated five scenarios for obtaining the genotype reference list of SNPs used in the demultiplexing algorithm: (i) list of population SNPs from the 1000 Genomes Project [21] provided by the authors of cellSNP/Vireo; (ii) list of population SNPs from the 1000 Genomes Project with an additional filtering step to retain only SNPs in the 3’ untranslated regions (UTRs) for faster runtime (this strategy is appropriate for 3’-tag sequencing protocols, but could also be adapted for 5’-tag or full-transcript sequencing); (iii) sample genotyping from matched bulk RNA-seq samples using bcftools [36]; (iv) sample genotyping from matched bulk RNA-seq samples using cellSNP [37]; and (v) sample genotyping from scRNA-seq samples using cellSNP [37]. Scenario (ii) was used for both datasets (HGSOC and lung adenocarcinoma), and the remaining scenarios were applied to the HGSOC dataset only. Scenarios (iii) and (iv) require matched bulk RNA-seq samples, while scenarios (i) and (v) have slow runtimes. Specifically, for the HGSOC dataset, we evaluated performance across several combinations of methods for genotyping and demultiplexing (labeled as “genotypeMethod_demultiplexingMethod” in Results). For the lung adenocarcinoma dataset, we used the list of population SNPs from the 1000 Genomes Project provided by the authors of cellSNP/Vireo, filtered to retain only SNPs in the 3’ UTRs.

For the main benchmark evaluations, we used two cancer datasets. The first dataset consists of three unique molecular identifier (UMI)-based scRNA-seq HGSOC samples measured on the 10x Genomics platform [43], obtained from separate, unrelated individuals at the Huntsman Cancer Institute at the University of Utah. We also obtained matched bulk RNA-seq samples from the same three individuals for sample genotyping. The raw data is available by controlled access via the Database of Genotypes and Phenotypes (dbGaP) (phs002262.v1.p1), and processed gene count tables are available from the Gene Expression Omnibus (GEO) (GSE158937). The second dataset consists of six UMI-based scRNA-seq higher-TMB lung adenocarcinoma samples measured on the 10x Genomics platform, previously published by [38]. Raw data for all samples in this study are available by controlled access from the European Genome-phenome Archive (EGA) (EGAD00001005054). For our study, we used six samples identified as having TMB >25 mutations / Mb (see [38], Figure 2d and Methods). **Table 1** and **Supplementary Table 2** provide a summary of the scRNA-seq datasets.

Performance was evaluated in terms of precision and recall for demultiplexing each scRNA-seq sample. We also recorded computational runtime for each step in the workflow and benchmark scenarios. Recall is defined as the proportion of true singlet cells for each sample that are identified as singlets and assigned to the correct sample. Precision is defined as the proportion of identified cells for each sample that are true singlet cells from the correct sample. Runtime was evaluated using the Unix date command. We used R version 4.1 for random number generation and evaluation steps performed in R, and created figures using ggplot2 [44].

For our benchmark evaluation, we developed three *in silico* simulation scenarios for each dataset -- containing either no doublets, 20% simulated doublets, or 30% simulated doublets. Doublets were simulated by combining cell barcode labels from random sets of two cells in the raw sequencing reads mapped using Cell Ranger [34], so that either 20% or 30% of the final barcodes represent doublets. For example, starting with 15,202 original cells in the HGSOC dataset, 3,508 randomly selected cells were combined with 3,508 other cells to create simulated doublets, leaving 11,694 final cell barcodes, of which 3,508 (30%) represent doublets. The 30% doublets scenario represents the upper end of our planned strategy for a “super-loading” experimental design, i.e. loading multiplexed cells at extremely high concentration to reduce library preparation costs and subsequently removing identifiable doublets [5,17,29]; the 20% doublets scenario represents an intermediate super-loading scenario; and the no doublets scenario serves as a best-case baseline scenario to evaluate performance of the demultiplexing tools.

### Ambient RNA from simulated cell debris

For the scenarios containing ambient RNA from simulated cell debris or lysed cells, we selected a percentage of cell barcodes (10%, 20%, or 40%) after doublet creation, and assigned all sequencing reads from these cell barcodes to other randomly selected cell barcodes, in each of the no doublets, 20% doublets, and 30% doublets scenarios. The debris percentages (10%, 20%, and 40%) were selected to be in the higher range of previously published results for non-cancer data [19], since ambient RNA is expected to be relatively abundant in complex or necrotic samples [40] such as cancer.

### Subset of SNPs from SNP array

For the simulated SNP array analyses, we selected a widely used SNP array (Infinium Multi-Ethnic Global-8 v1.0 array from the Multi-Ethnic Genotyping Array Consortium (MEGA) Consortium), and calculated the overlapping sets of SNPs between the total 1.7 million SNPs from the array and our existing genotype references (see **Supplementary Table 1** for a summary of the overlapping set sizes). Then, we re-ran our benchmark evaluations using the subsets of SNPs from the overlaps as the genotype references.

### Healthy (non-cancer) cell line data

For a baseline comparison with healthy (non-cancer) data, we combined sequencing reads from 5 samples of induced pluripotent stem cell (iPSC) cell lines from the Human iPSC Initiative (HipSci), which were previously published by [19], and added the same percentages of doublets (no doublets, 20% doublets, or 30% doublets) as in our main analyses. The raw data are available from the European Nucleotide Archive (ENA) (ERS2630502-ERS2630506). Compared to our cancer samples, these samples contained relatively higher numbers of cells per sample, as well as higher unique molecular identifier (UMI) counts per cell (details are provided in **Supplementary Table 2**).

### Single-cell RNA sequencing of ovarian tumors

De-identified HGSOC samples were processed after cryopreservation in liquid nitrogen where tissue chunks were stored in RPMI media with 10% fetal bovine serum and 10% DMSO. Samples were thawed and dissociated to single cells using the Miltenyi Human Tumor Dissociation Kit and the GentleMACS dissociator. Samples were incubated on the GentleMACS at 37°C for 1 hour with the setting of 1,865 rounds per run. A 70 µm MACS smart strainer was used to deplete cell doublets before loading onto the 10x Genomics Chromium Controller. Library preparation was performed using the 10x Genomics 3’ Gene Expression Library Prep v3 and libraries were sequenced on an Illumina NovaSeq instrument.

## Availability of source code and requirements

Project name: snp-dmx-cancer

Project home page: https://github.com/lmweber/snp-dmx-cancer

Operating system: Linux

Programming language: Shell, R

Other requirements: High performance computing (HPC) cluster with Sun Grid Engine (SGE) scheduler

License: MIT

## Code availability

All code scripts to reproduce the benchmark evaluations, supplementary analyses, generate figures in the manuscript, and run the Snakemake workflow are freely accessible from GitHub at https://github.com/lmweber/snp-dmx-cancer. All tools used within the benchmark evaluations and workflow are freely available, as described in Methods. Software versions used were Cell Ranger 4.0.0, bcftools 1.10.2-91-g365d117, demuxlet 3ab507c, cellsnp-lite 1.2.0, and Vireo 0.5.0.

## Data availability

Raw and processed sequencing data generated in this study (HGSOC dataset) are available from the Database of Genotypes and Phenotypes (dbGaP) (raw data consisting of FASTQ files, accession phs002262.v1.p1) and Gene Expression Omnibus (GEO) (processed data files containing gene count tables, accession GSE158937). The lung adenocarcinoma dataset was previously published by [36], and is available from the European Genome-phenome Archive (EGA) (EGAD00001005054). The healthy (non-cancer) iPSC cell line data were previously published by [19], and are available from the European Nucleotide Archive (ENA) (ERS2630502-ERS2630506).

## Acknowledgments

We thank Yuanhua Huang for assistance with running Vireo and cellSNP; Davis McCarthy for advice regarding Vireo; and attendees from the Stephanie Hicks and Kasper Hansen joint lab meetings at Johns Hopkins University for helpful feedback and discussions. We thank Hae-Ock Lee and Myung-Ju Ahn of the Samsung Medical Center for providing access to the lung adenocarcinoma dataset. Research reported in this publication utilized the Biorepository and Molecular Pathology Shared Resource and the High-Throughput Genomics Shared Resource at the Huntsman Cancer Institute at University of Utah and was supported by NIH/NCI award P30 CA042014. Computational analyses were performed using the Joint High Performance Computing Exchange (JHPCE) high-performance computing facility in the Department of Biostatistics at the Johns Hopkins Bloomberg School of Public Health. The content is solely the responsibility of the authors and does not necessarily represent the official views of the NIH.

## Author contributions

LMW: Software, Formal analysis, Investigation, Data curation, Writing - Original Draft, Visualization

AAH: Software

PFH: Software, Writing - Review & Editing

KCB: Investigation

JG: Investigation, Resources, Writing - Review & Editing

JAD: Resources, Writing - Review & Editing, Funding acquisition

CSG: Conceptualization, Resources, Writing - Review & Editing, Funding acquisition

SCH: Conceptualization, Resources, Writing - Original Draft, Writing - Review & Editing, Supervision, Funding acquisition

## Ethics approval and consent to participate

Ovarian cancer tissue was obtained and studied under written informed consent at the Huntsman Cancer Institute through approved University of Utah Institutional Review Board protocols IRB_00010924 and IRB_00118086. Analysis of human data in this study was also approved by the University of Pennsylvania Institutional Review Board (IRB protocol 832353) and the Johns Hopkins Bloomberg School of Public Health Institutional Review Board (IRB00013099).

## Supplementary Figures

**Supplementary Figure 1.**
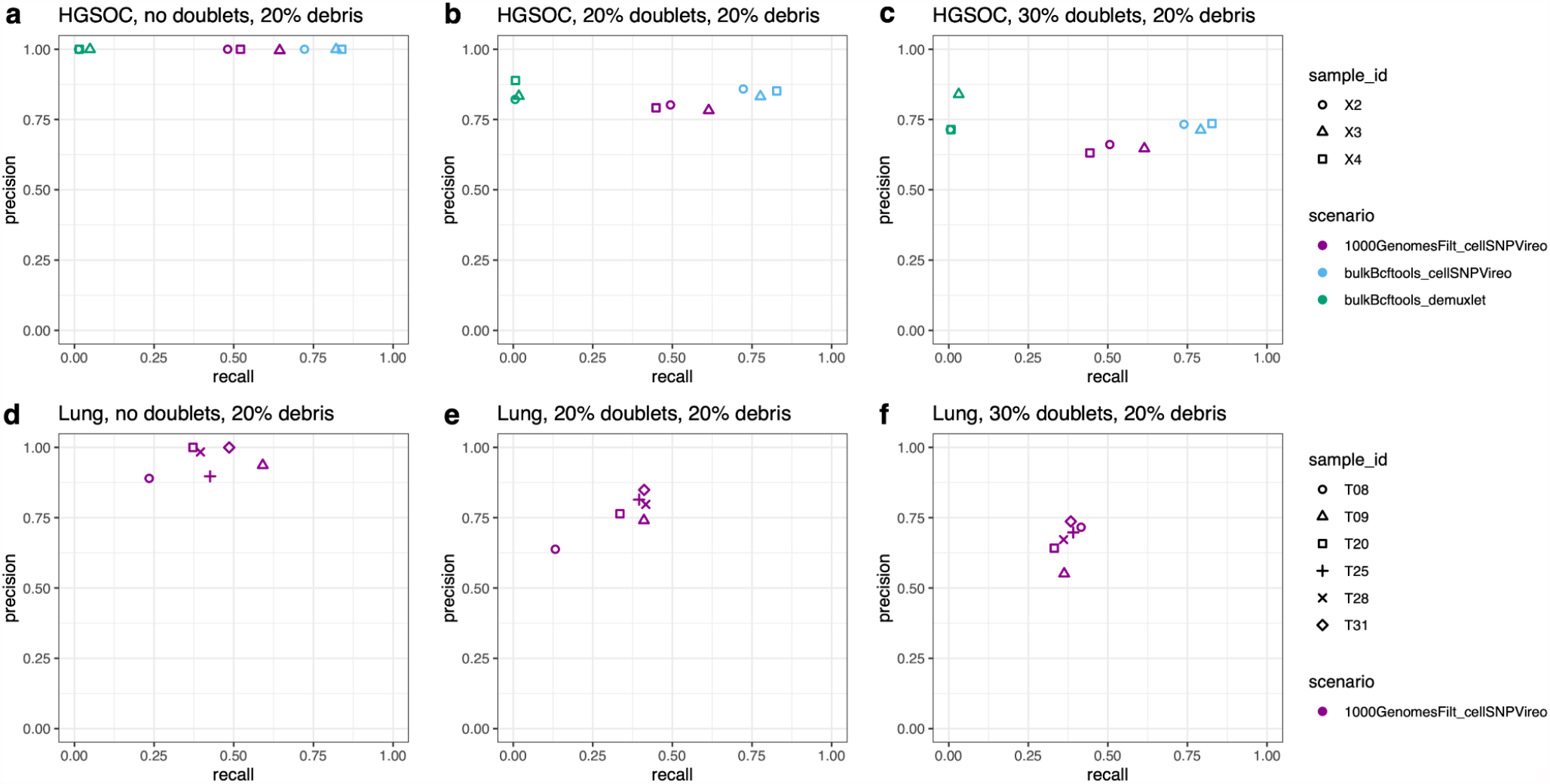
Performance evaluations for benchmark scenarios including ambient RNA from simulated cell debris. Top-performing and computationally efficient scenarios for HGSOC dataset **(a-c)** and lung adenocarcinoma dataset **(d-f)**, across three proportions of simulated doublets (no doublets, 20% doublets, 30% doublets), after introducing ambient RNA from simulated cell debris by assigning all reads from 20% of final cell barcodes to other randomly selected cell barcodes. Performance is evaluated in terms of precision (y-axis) and recall (x-axis) for recovering the sample identities of true singlet cells from each scRNA-seq sample. Benchmark scenarios are labeled by color and with the naming scheme “genotypeMethod_demultiplexingMethod”. Samples within each dataset are identified with shapes. Axis limits range from 0 to 1 for all panels

**Supplementary Figure 2.**
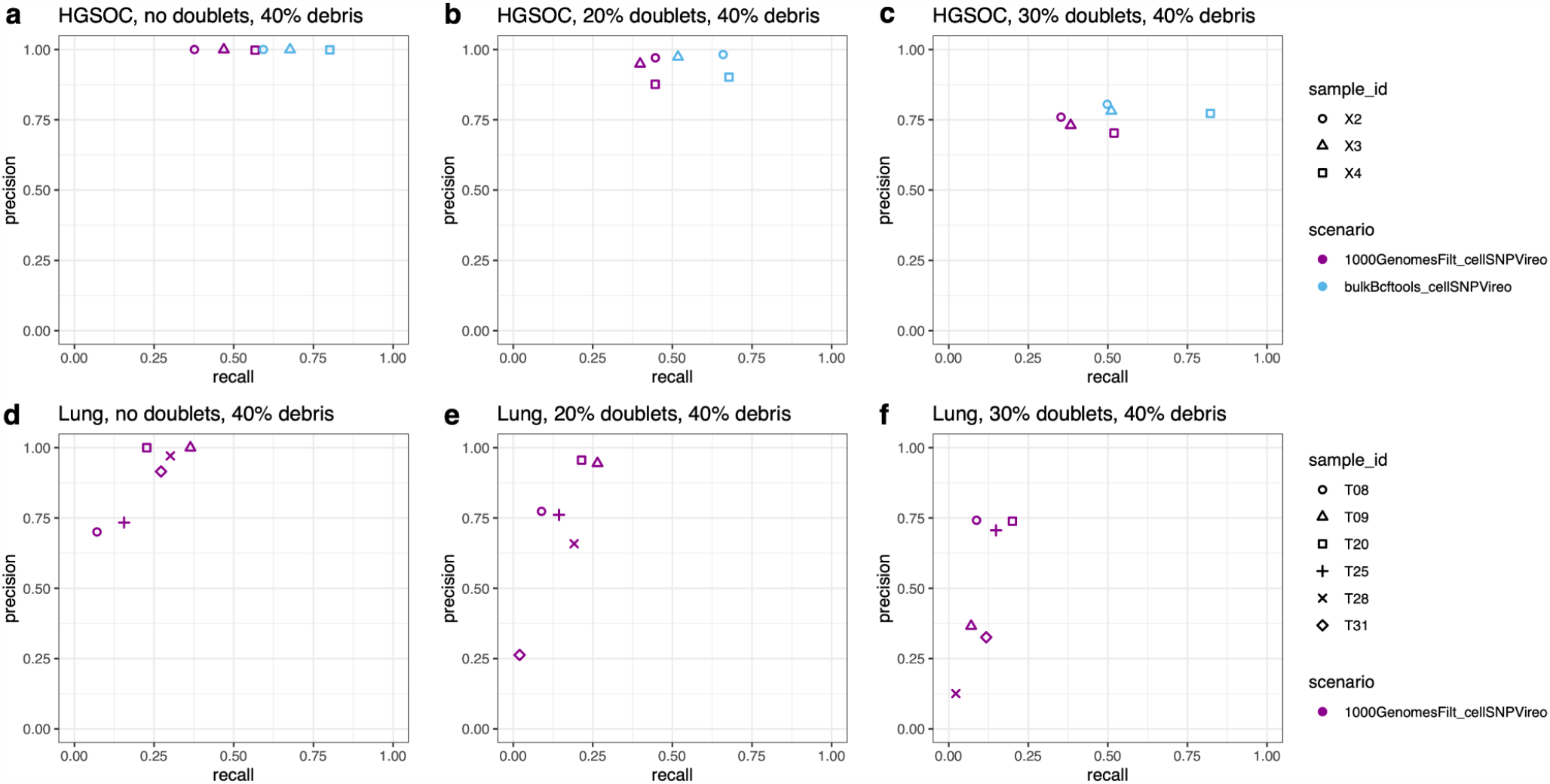
Performance evaluations for benchmark scenarios including ambient RNA from simulated cell debris. Top-performing and computationally efficient scenarios for HGSOC dataset **(a-c)** and lung adenocarcinoma dataset **(d-f)**, across three proportions of simulated doublets (no doublets, 20% doublets, 30% doublets), after introducing ambient RNA from simulated cell debris by assigning all reads from 40% of final cell barcodes to other randomly selected cell barcodes. Note that unlike Figure 3 and Supplementary Figure 1, demuxlet is not included for the HGSOC dataset, since this tool did not successfully run with this higher proportion of ambient RNA. Performance is evaluated in terms of precision (y-axis) and recall (x-axis) for recovering the sample identities of true singlet cells from each scRNA-seq sample. Benchmark scenarios are labeled by color and with the naming scheme “genotypeMethod_demultiplexingMethod”. Samples within each dataset are identified with shapes. Axis limits range from 0 to 1 for all panels.

**Supplementary Figure 3.**
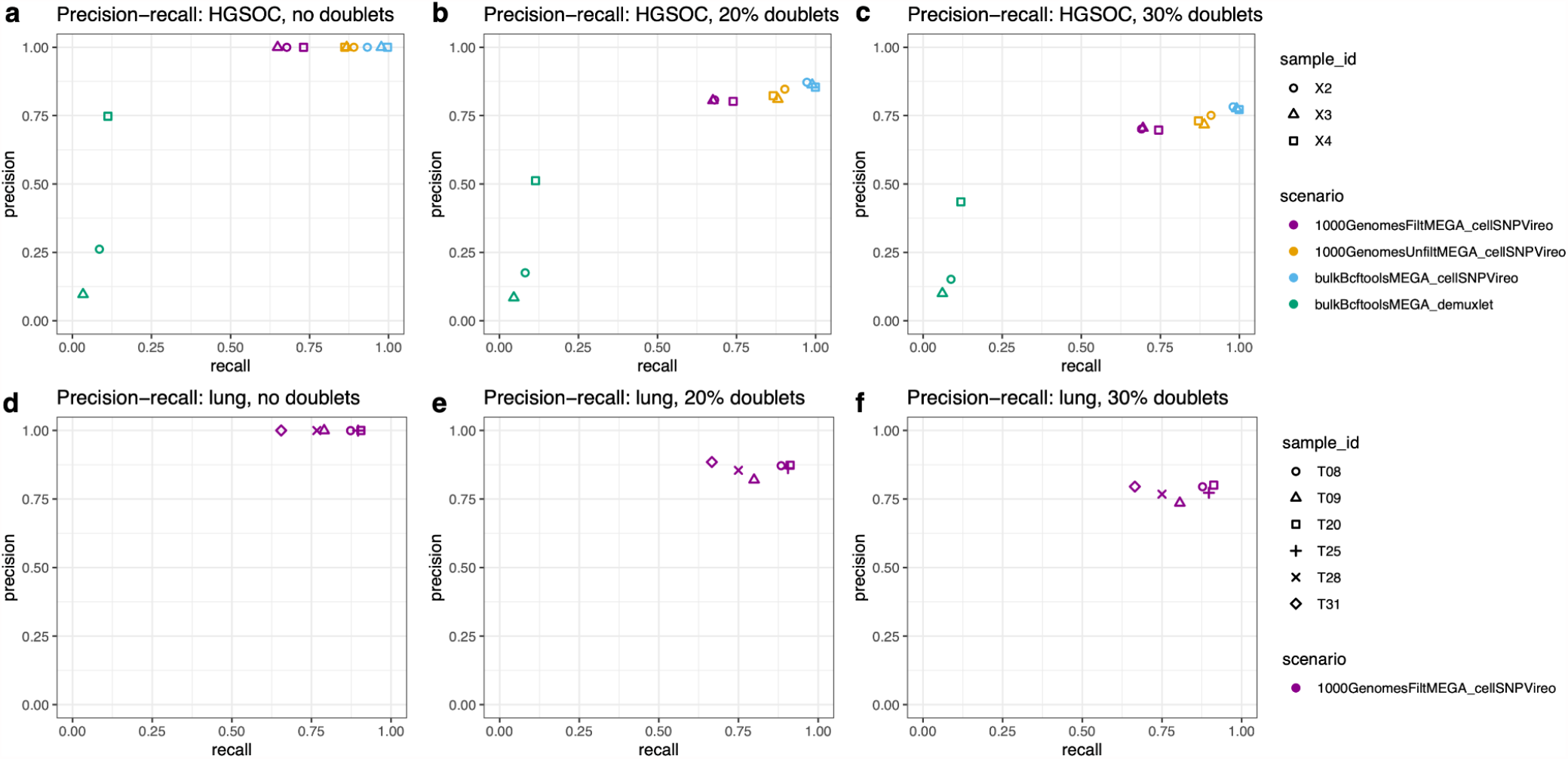
Performance evaluations for benchmark scenarios using subset of SNPs from SNP array as genotype reference. Top-performing and computationally efficient scenarios for HGSOC dataset **(a-c)** and lung adenocarcinoma dataset **(d-f)**, across three proportions of simulated doublets (no doublets, 20% doublets, 30% doublets), when using a subset of SNPs from a SNP array (Infinium Multi-Ethnic Global-8 v1.0 array from the Multi-Ethnic Genotyping Array Consortium (MEGA) Consortium) overlapping with either the 1000 Genomes filtered (“1000GenomesFiltMEGA”), 1000 Genomes unfiltered (“1000GenomesUnfiltMEGA”), or bulk RNA-seq (“bulkBcftoolsMEGA”) reference as the genotype reference for demultiplexing. Performance is evaluated in terms of precision (y-axis) and recall (x-axis) for recovering the sample identities of true singlet cells from each scRNA-seq sample. Benchmark scenarios are labeled by color and with the naming scheme “genotypeMethod_demultiplexingMethod”. Samples within each dataset are identified with shapes. Axis limits range from 0 to 1 for all panels.

**Supplementary Figure 4.**
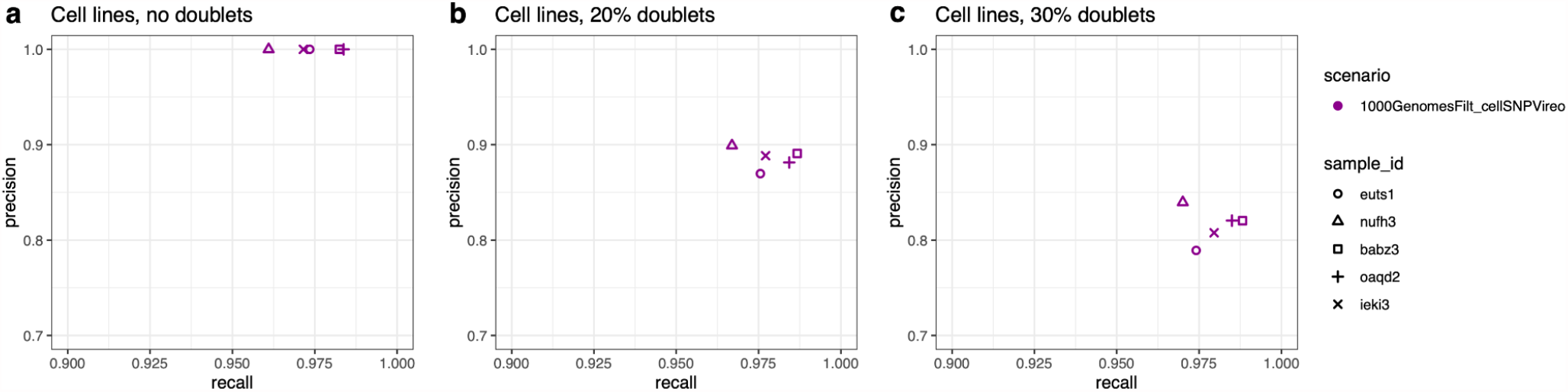
Performance evaluations for healthy (non-cancer) cell line dataset. Top-performing and computationally efficient scenario from main results (“1000GenomesFilt_cellSNPVireo”) for healthy (non-cancer) induced pluripotent stem cell (iPSC) cell line dataset sourced from [19], consisting of 5 samples, across three proportions of simulated doublets **(a-c)** (no doublets, 20% doublets, and 30% doublets). Performance is evaluated in terms of precision (y-axis) and recall (x-axis) for recovering the sample identities of true singlet cells from each scRNA-seq sample. Benchmark scenarios are labeled by color and with the naming scheme “genotypeMethod_demultiplexingMethod”. Samples within each dataset are identified with shapes. Axis limits differ between y-axis and x-axis for improved visibility, and are the same in all panels.

**Supplementary Figure 5.**
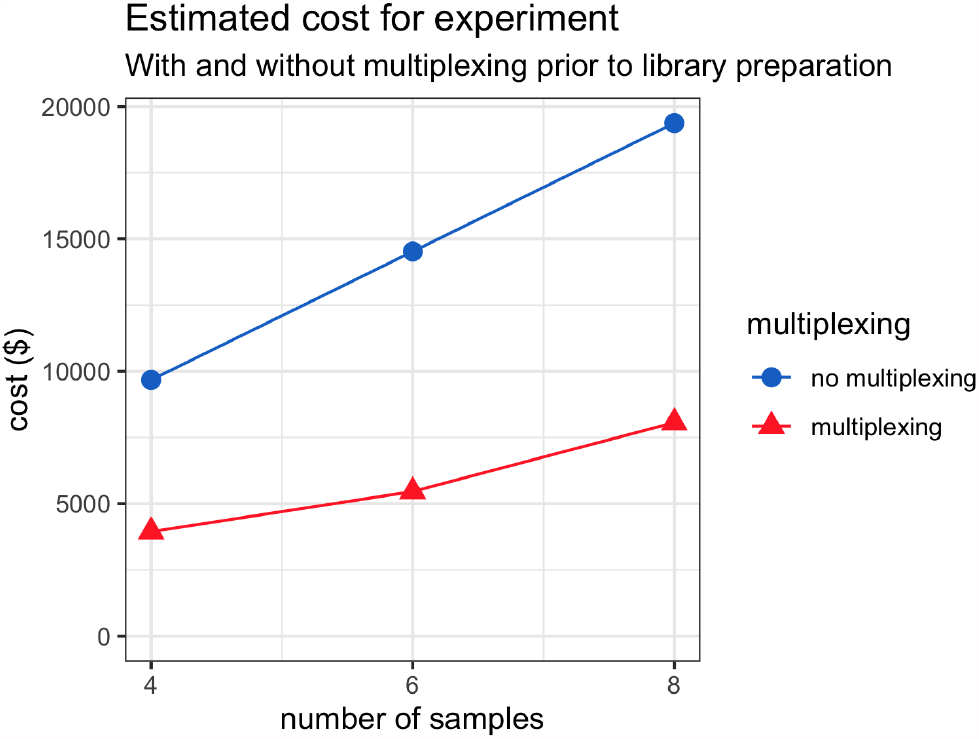
Illustration of expected cost savings from multiplexed experimental design prior to library preparation. The figure shows the total of estimated library preparation and sequencing costs, with either no multiplexing or full multiplexing (all samples prepared as a single library and sequenced together), for experiments with 4, 6, or 8 samples. The calculations assume 4,000 desired cells per sample after demultiplexing, after discarding identifiable doublets consisting of cells from multiple samples; library preparation costs of $2,000 per sample or multiplexed set of samples; sequencing costs of $1,500 per 400 million reads with an additional 30% cost due to unaligned reads and adapters; and approximately 20,000 reads per cell. Calculations were performed using the “Cost Per Cell” online calculator provided by the Satija Lab [29].

## Supplementary Tables

**Supplementary Table 1.** Number and percentage of SNPs in sets overlapping between the available genotype references (bulk RNA-seq from 3 HGSOC samples, 1000 Genomes filtered to 3’ UTRs, 1000 Genomes unfiltered, and MEGA SNP array). The bulk RNA-seq and 1000 Genomes filtered references are used for the main results.

|  | bulk RNA-seq<br>(3 HGSOC<br>samples) | 1000 Genomes<br>(filtered 3' UTRs) | 1000 Genomes<br>(unfiltered) | SNP array (MEGA) |
| --- | --- | --- | --- | --- |
| bulk RNA-seq (3<br>HGSOC samples) | 605,367 | 28,501 | 356,812 | 45,843<br>(7.6% of bulk RNA-seq)<br>(2.6% of SNP array) |
| 1000 Genomes<br>(filtered 3' UTRs) |  | 84,853 | 84,853 | 14,021<br>(16.5% of 1000 Genomes filtered)<br>(0.8% of SNP array) |
| 1000 Genomes<br>(unfiltered) |  |  | 7,414,539 | 638,657<br>(8.6% of 1000 Genomes unfiltered)<br>(36.8% of SNP array) |
| SNP array (MEGA) |  |  |  | 1,733,345 |

**Supplementary Table 2.**
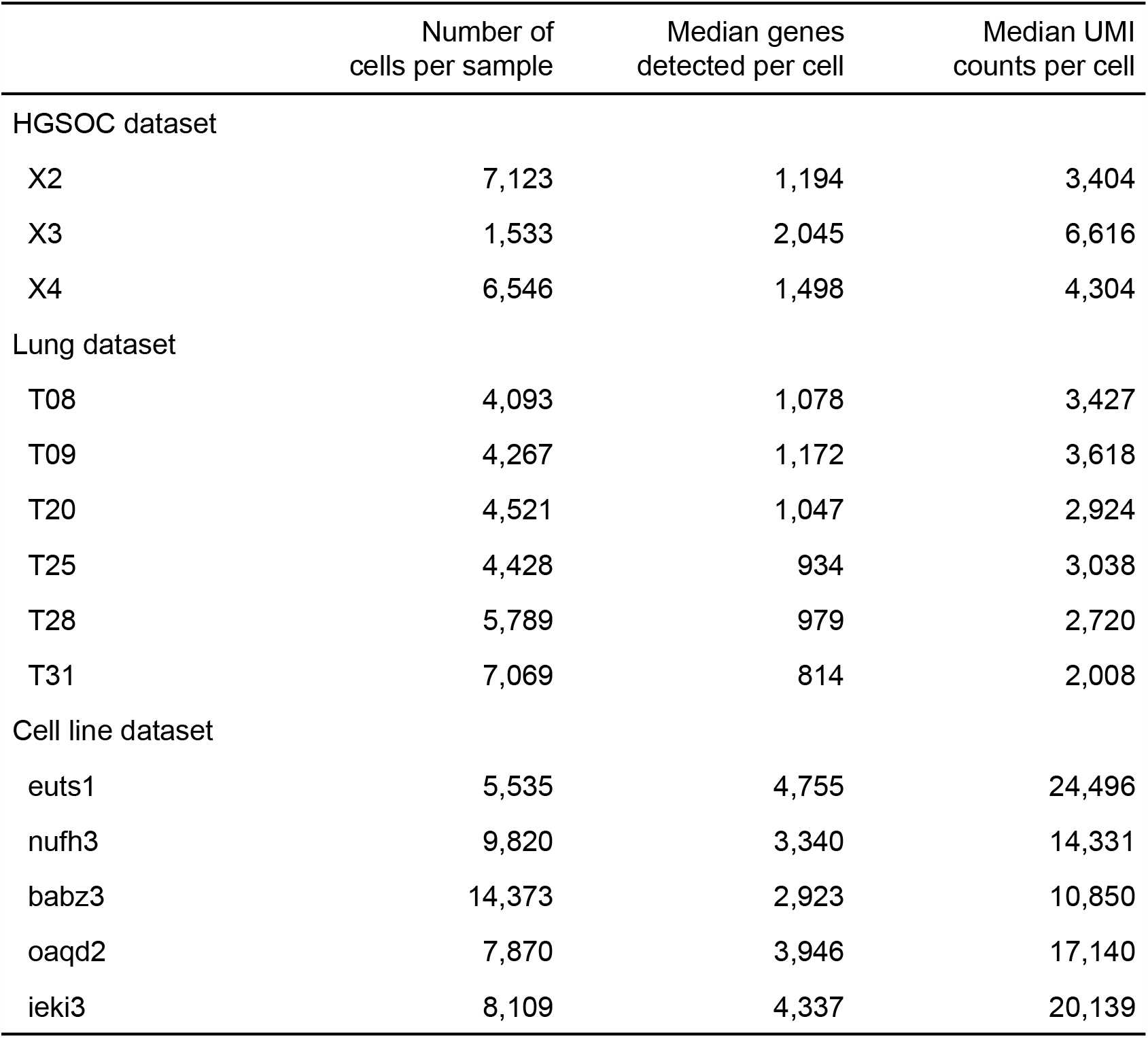
Summary of HGSOC, lung adenocarcinoma [38], and healthy induced pluripotent stem cells (iPSC) cell line [19] datasets. Number of cells per sample, median genes detected per cell, and median unique molecular identifier (UMI) counts per cell are shown for each dataset. Median genes detected and median UMI counts per cell were higher for the cell line dataset than for the HGSOC and lung adenocarcinoma datasets.

**Supplementary Table 3.** Confusion matrix for singlet, doublet, and unassigned calls for the top-performing scenario (cellSNP/Vireo with bulk RNA-seq genotype reference) for 30% doublets scenario for HGSOC dataset (matching the precision-recall values in the main results shown in **Figure 2 c**). Calls by Vireo are shown in columns (singlets: donor0, donor1, donor2 in arbitrary sample order; doublets; unassigned), and true labels from the simulation are shown in rows (singlets: X2, X3, X4; non-identifiable doublets from the same sample: X2-X2, X3-X3, X4-X4; identifiable doublets: dbl-X2-X3, dbl-X2-X4, dbl-X3-X4). Doublets consisting of two cells from the same sample are non-identifiable since these cells contain the same germline SNPs.

| HGSOC, 30% doublets, bulkBcftools_cellSNPVireo |  |  |  |  |  |
| --- | --- | --- | --- | --- | --- |
|  | donor1 | donor0 | donor2 | doublet | unassigned |
| X2 | 3839 | 0 | 8 | 7 | 1 |
| X2-X2 | 757 | 0 | 1 | 2 | 0 |
| X3 | 0 | 826 | 0 | 0 | 0 |
| X3-X3 | 0 | 38 | 0 | 0 | 0 |
| X4 | 0 | 0 | 3505 | 0 | 0 |
| X4-X4 | 0 | 0 | 647 | 0 | 0 |
| dbl-X2-X3 | 46 | 107 | 0 | 162 | 1 |
| dbl-X2-X4 | 223 | 0 | 343 | 859 | 7 |
| dbl-X3-X4 | 0 | 114 | 41 | 160 | 0 |
| % true identifiable doublets | 5.5% | 20.4% | 8.4% | 99.2% | 88.9% |

**Supplementary Table 4.** Confusion matrix for singlet, doublet, and ambiguous calls for demuxlet (with bulk RNA-seq genotype reference) for 30% doublets scenario for HGSOC dataset (matching the precision-recall values in the main results shown in **Figure 2 c**). Calls by demuxlet are shown in columns (singlets: X2, X3, X4; doublets; ambiguous), and true labels from the simulation are shown in rows (singlets: X2, X3, X4; non-identifiable doublets from the same sample: X2-X2, X3-X3, X4-X4; identifiable doublets: dbl-X2-X3, dbl-X2-X4, dbl-X3-X4). Doublets consisting of two cells from the same sample are non-identifiable since these cells contain the same germline SNPs

| HGSOC, 30% doublets, bulkBcftools_demuxlet |  |  |  |  |  |
| --- | --- | --- | --- | --- | --- |
|  | X2 | X3 | X4 | doublet | ambiguous |
| X2 | 2560 | 0 | 0 | 993 | 0 |
| X2-X2 | 510 | 0 | 0 | 221 | 0 |
| X3 | 0 | 340 | 0 | 486 | 0 |
| X3-X3 | 0 | 13 | 0 | 25 | 0 |
| X4 | 0 | 1 | 1712 | 1776 | 3 |
| X4-X4 | 0 | 0 | 302 | 345 | 0 |
| dbl-X2-X3 | 15 | 14 | 0 | 287 | 0 |
| dbl-X2-X4 | 38 | 0 | 40 | 1225 | 0 |
| dbl-X3-X4 | 0 | 19 | 7 | 289 | 0 |
| % true identifiable doublets | 1.7% | 8.5% | 2.3% | 31.9% | 0.0% |

**Supplementary Table 5.** Summary of doublets identified by applying a downstream doublet detection tool (scDblFinder) to demultiplexed cells after applying top-performing and computationally efficient demultiplexing tools (cellSNP/Vireo with bulk RNA-seq reference for HGSOC dataset; cellSNP/Vireo with 1000 Genomes 3’ UTRs filtered reference for lung adenocarcinoma dataset), for 20% and 30% doublets scenarios. scDblFinder was run using default settings, and clusters representing doublets identified using thresholds of 70 (HGSOC) and 10 (lung adenocarcinoma) differentially expressed genes per cluster based on inspection of elbow plots. False positive and false negative doublet calls are shown in bold font.

| HGSOC, 20% doublets |  |  | HGSOC, 30% doublets |  |  |
| --- | --- | --- | --- | --- | --- |
| identified as doublet |  |  | identified as doublet |  |  |
| true doublet | TRUE | FALSE | true doublet | TRUE | FALSE |
| TRUE | 600 | **1,003** | TRUE | 917 | **1,408** |
| FALSE | **3,720** | 6,405 | FALSE | **3,214** | 4,965 |

**Supplementary Table 5.** Summary of doublets identified by applying a downstream doublet detection tool (scDbIFinder) to demultiplexed cells after applying top-performing and computationally efficient demultiplexing tools (cellSNP/Vireo with bulk RNA-seq reference for HGSOC dataset; cellSNP/Vireo with 1000 Genomes 3' UTRs filtered reference for lung adenocarcinoma dataset), for 20% and 30% doublets scenarios.
| lung, 20% doublets |  |  | lung, 30% doublets |  |  |
| --- | --- | --- | --- | --- | --- |
| identified as doublet |  |  | identified as doublet |  |  |
| true doublet | TRUE | FALSE | true doublet | TRUE | FALSE |
| TRUE | 346 | **2,221** | TRUE | 326 | **3,328** |
| FALSE | **2,088** | 17,678 | FALSE | **1,129** | 14,798 |

